# Adiponectin pathway activation dampens inflammation and enhances alveolar macrophage fungal killing via LC3-associated phagocytosis

**DOI:** 10.1101/2024.06.24.600373

**Authors:** Sri Harshini Goli, Joo-Yeon Lim, Nese Basaran-Akgul, Steven P. Templeton

## Abstract

Although innate immunity is critical for antifungal host defense against the human opportunistic fungal pathogen *Aspergillus fumigatus*, potentially damaging inflammation must be controlled. Adiponectin (APN) is an adipokine produced mainly in adipose tissue that exerts anti-inflammatory effects in adipose-distal tissues such as the lung. We observed 100% mortality and increased fungal burden and inflammation in neutropenic mice with invasive aspergillosis (IA) that lack APN or the APN receptors AdipoR1 or AdipoR2. Alveolar macrophages (AMs), early immune sentinels that detect and respond to lung infection, express both receptors, and APN-/- AMs exhibited an inflammatory/M1 phenotype that was associated with decreased fungal killing. Pharmacological stimulation of AMs with AdipoR agonist AdipoRon partially rescued deficient killing in APN-/- AMs that was dependent on both receptors. Finally, APN-enhanced fungal killing was associated with increased activation of the non-canonical LC3 pathway of autophagy. Thus, our study identifies a novel role for APN in LC3-mediated killing of *A.fumigatus*.

**Author Summary:** *Aspergillus fumigatus* is a human fungal pathogen that causes an often-fatal invasive infection of the lung in immune compromised individuals, a population that is increasing. Since current antifungal drugs have limited efficacy, it is important to identify pathways that may be pharmaceutically targeted to complement existing therapies. Adiponectin (APN) is an anti-inflammatory intercellular cytokine messenger produced mainly in adipose tissue that protects against invasive aspergillosis. Alveolar macrophages are early immune sentinel cells in the lung, and we report that AMs from mice lacking APN exhibit an inflammatory phenotype and reduced killing of *A. fumigatus* spores that is improved when AMs are treated with a drug (AdipoRon) that simulates APN binding to its receptors AdipoR1 and AdipoR2. Furthermore, APN was associated with activation of LC3-associated phagocytosis, a mechanism of fungal killing important for host defense against *A. fumigatus* infection. Thus, we identify therapeutic potential of the APN pathway in stimulation of immune-mediated fungal killing and treatment of fungal infection.

## Introduction

The human opportunistic fungal pathogen *Aspergillus fumigatus* is the primary etiologic agent of invasive pulmonary aspergillosis (IA), a frequently severe infection with a high mortality rate [1]. Effective treatment options for IA remain limited, despite increases in the at-risk population, and fungal resistance to existing antifungal drugs is increasing. Furthermore, although an underlying immune deficiency renders individuals susceptible to IA, poor disease outcomes are associated with detrimental inflammatory pathology [2,3]. It is thus critical to identify anti-inflammatory pathways that could be exploited to target detrimental immune pathology, enhance antifungal immunity, and complement existing antifungal therapies in IA patients.

Adiponectin (APN) is an anti-inflammatory adipokine produced mainly in adipose tissue and expressed at high levels in lean and healthy individuals, whereas in obese individuals, adipose tissue is inflamed and APN production and secretion are inhibited [4–6]. Numerous studies have demonstrated a protective effect for APN in autoimmune and inflammatory diseases, and immune-mediated clearance of *Listeria monocytogenes* was hampered in obese and APN-deficient mice due to detrimental inflammation and hematopoietic dysfunction [7]. More recently, we demonstrated increased mortality, fungal burden, lung inflammatory cytokine production and eosinophil recruitment in APN-deficient mice with IA [8]. However, eosinophils did not play a significant role in the pathology of APN-deficiency, as neutropenic mice deficient in both APN and eosinophils exhibited only delayed, but not decreased mortality, suggesting other unidentified mechanisms of APN-mediated protection in IA.

Alveolar macrophages (AMs) are the primary immune sentinel of lung airways, and their early responses to inhaled fungal conidia are critical for disease outcomes, particularly in immunocompromised hosts [9,10]. In our previous study, we observed increased production of the pro-inflammatory cytokine TNF in APN-/- AMs [8], suggesting that these cells contribute to the inflammatory phenotype of APN-/- mice with IA. In this study, we aimed to determine how the adiponectin pathway regulates responses of AMs to *A. fumigatus* infection. We report increased production of inflammatory cytokines, an elevated M1 phenotype, and reduced uptake and killing of conidia in APN-/- AMs that is increased by stimulation with the APN receptor agonist AdipoRon. Furthermore, APN stimulation was associated with increased AM activation of the non-canonical autophagy LC3 pathway in conidia-containing phagosomes, thus identifying a novel role for APN in promoting LC3-associated phagocytosis (LAP) in antifungal immunity.

## Results

### Increased mortality, fungal burden, and inflammatory pathology in APN pathway-deficient mice with IA

Previously, we reported that APN inhibits inflammatory lung pathology during IA [8]. Since APN-/- strains vary in their metabolic phenotypes[11–13], we confirmed the inflammatory phenotype of APN-/- mice from a second strain (obtained from Dr. Philipp Scherer) (**Fig. S1**), and observed similar increases in fungal burden (**Figs. S1B-S1D**), inflammatory cytokines and inflammatory cell recruitment (**Figs. S1E-S1F** and **S1H**) while only survival and AdipoR1 transcription were not significantly different from wild-type (WT) C57BL/6 controls (**Figs. S1A** and **S1G**, respectively). We also aimed to determine the contribution of the canonical adiponectin receptors AdipoR1 and AdipoR2 to IA pathology, and thus infected neutrophil-depleted WT, APN-/-, AdipoR1-/- and AdipoR2-/- mice to compare mortality, pathology, fungal burden, inflammatory cell recruitment and cytokine production (**Fig. 1A**). In contrast to WT mice, mortality was 100% in APN-/-, AdipoR1-/-, and AdipoR2-/- mice, with mortality beginning on day 2 post-infection and 100% by day 6 p.i. (**Fig. 1B**), with extensive lung hyphal growth and peri-bronchoalveolar inflammation (**Fig. 1C**) and increased fungal burden as measured by qPCR of fungal DNA (**Fig. 1D**) and image quantification of fungal GMS staining (**Fig. 1E**). Inflammatory cells were increased in the bronchoalveolar lavage fluid (BALF), most significantly in APN-/- mice (**Fig. 1F**), while lung inflammatory cytokines IL-1a, IL-6, TNF, IL-12b, and IL-17A and the anti-inflammatory cytokine IL-10 were increased, most significantly in AdipoR1-/- mice (**Figs. 1G and 1H**). In summary, adiponectin-pathway deficiency is associated with increased mortality, lung fungal burden, and inflammatory pathology, cell recruitment, and cytokine production. Furthermore, disrupted signaling through either AdipoR1 or AdipoR2 results in increased IA severity.

**Figure 1.**
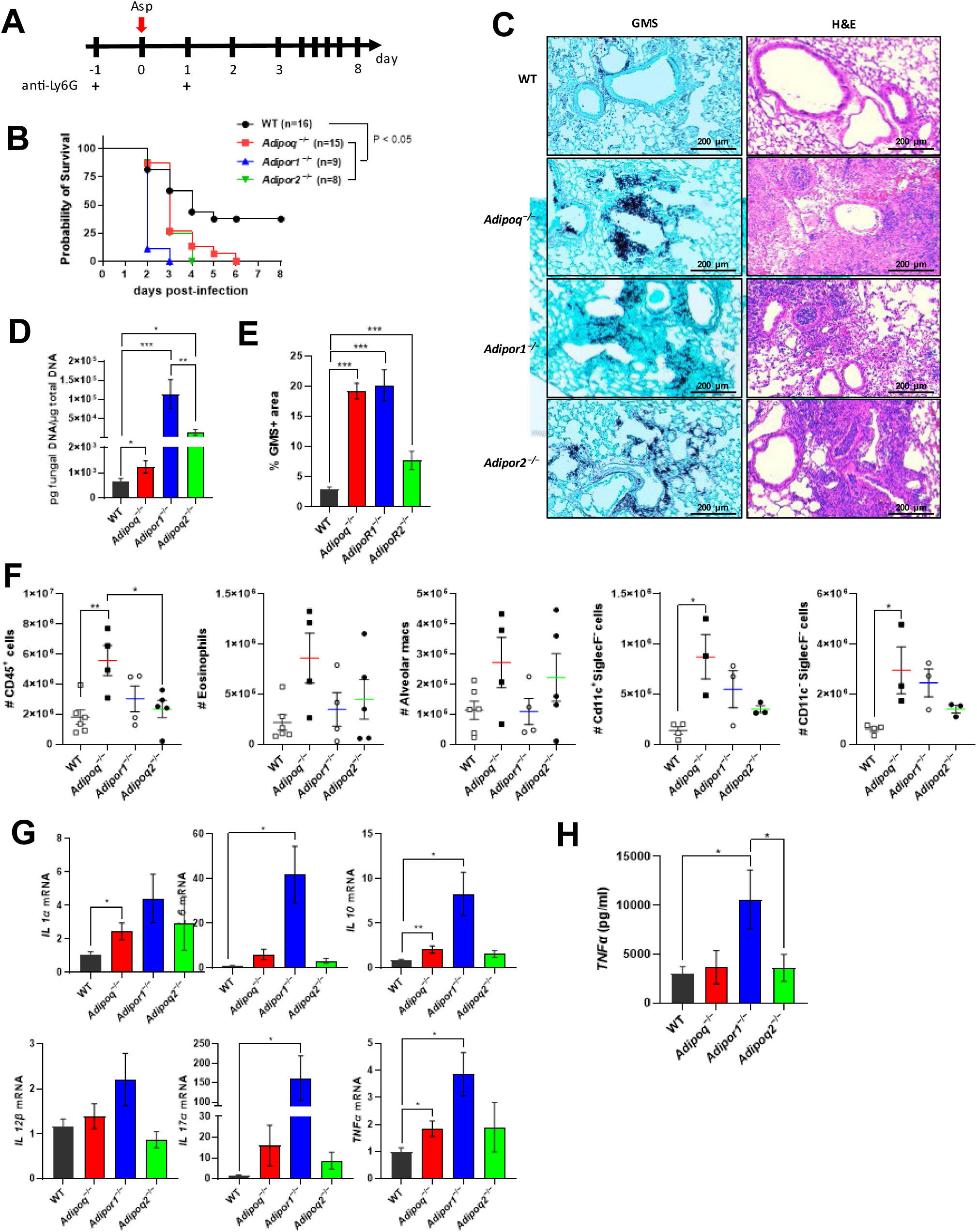
Increased mortality, fungal burden, and inflammation in APN pathway-deficient mice with invasive aspergillosis. Wild-type (C57BL/6), APN-/- (*Adipoq*^−/−^), *AdipoR1*^−/−^, and *AdipoR2*^−/−^ mice were neutrophil-depleted and involuntarily aspirated a suspension of *A. fumigatus* conidia and followed for survival or sacrificed on day 3 post-infection as described in Materials and Methods. A. Experimental time course for neutropenic mouse model of infection. B. Survival. *N* = 8-16 mice per group. C. Representative GMS and H&E lung sections. D. Fungal burden determined by quantitative PCR of fungal DNA from lung homogenates. E. Fungal burden determined by quantification of GMS staining. F. Total number of CD45^+^ cells, eosinophils (CD45^+^Ly6G^−^CD11c^−^SiglecF^+^), AMs (CD45^+^Ly6G^−^CD11c^+^SiglecF^+^), CD11c^+^SiglecF^−^ (CD45^+^Ly6G^−^ CD11c^+^SiglecF^−^), and CD11c^−^SiglecF^−^ (CD45^+^Ly6G^−^CD11c^−^SiglecF^−^) cells isolated from the mice with IA as determined by flow cytometry. *N* = 4-6 mice per group. G. qRT-PCR analysis for mRNA expression of the indicated cytokines. H. TNFα secretion in BALF quantified by ELISA. A-D, data are a summary of two to three independently performed experiments. F-H, data are representative of two independent experiments with similar results. \**p* < 0.05, \*\**p* < 0.01, \*\*\**p* < 0.001.

### Alveolar macrophages (AMs) from adiponectin pathway-deficient mice exhibit an inflammatory phenotype

Previously, we observed increased intracellular TNF in AMs from APN-/- mice with IA when compared to WT AMs [8], and we observed mortality as early as day 2 post-infection in APN-pathway-deficient mice with IA, suggesting a critical involvement of early immune effectors in protection from infection. As AMs contribute to early responses to inhalation of *A. fumigatus* [9], especially in immunocompromised hosts [10], we aimed to further examine the phenotype of AMs from APN pathway-deficient hosts using primary ex vivo cultures. We observed expression of both AdipoR1 and AdipoR2 in WT and APN-deficient mice that was decreased in lung tissue upon *A. fumigatus* infection, but increased in the BALF (**Fig. S2A**), with the highest surface expression of AdipoR1 in AMs (**Fig. S2B**). APN-pathway deficient AMs exhibited differential expression of AdipoRs at the transcription level **Fig. S2C**), and infection of APN-/- AMs decreased AdipoR1 expression (**Fig. S2D**). Although WT, APN-/-, AdipoR1-/-, and AdipoR2-/- AMs were similar morphologically (data not shown), transcription of IL-1α, IL-1β, IL-6, IL-17A, IL-23, and TNF was increased in APN pathway-deficient AMs when compared WT-derived AMs, while IL-10 was similar (**Fig. 2A**), and secreted TNF in the supernatant increased equally in response to infection in all groups (**Fig. 2B**). To investigate the inflammatory phenotype more specifically in ex vivo cultured APN-/- AMs, we compared expression of the macrophage polarization markers CD38 (M1/classical) and Egr2 (M2/alternative) with wild type WT AMs [14]. In a resting state, a higher percentage of APN-/- AMs were CD38+ and lower percentage were Egr2+ when compared to WT (**Figs. 2C and 2D**), while upon infection with *A. fumigatus*, percentages of both CD38+Egr2- and CD38-Egr2+ cells increased (**Figs. 2C and 2E**). These results suggest that APN pathway-deficient AMs exhibit an increased inflammatory phenotype compared to WT AMs.

**Figure 2.**
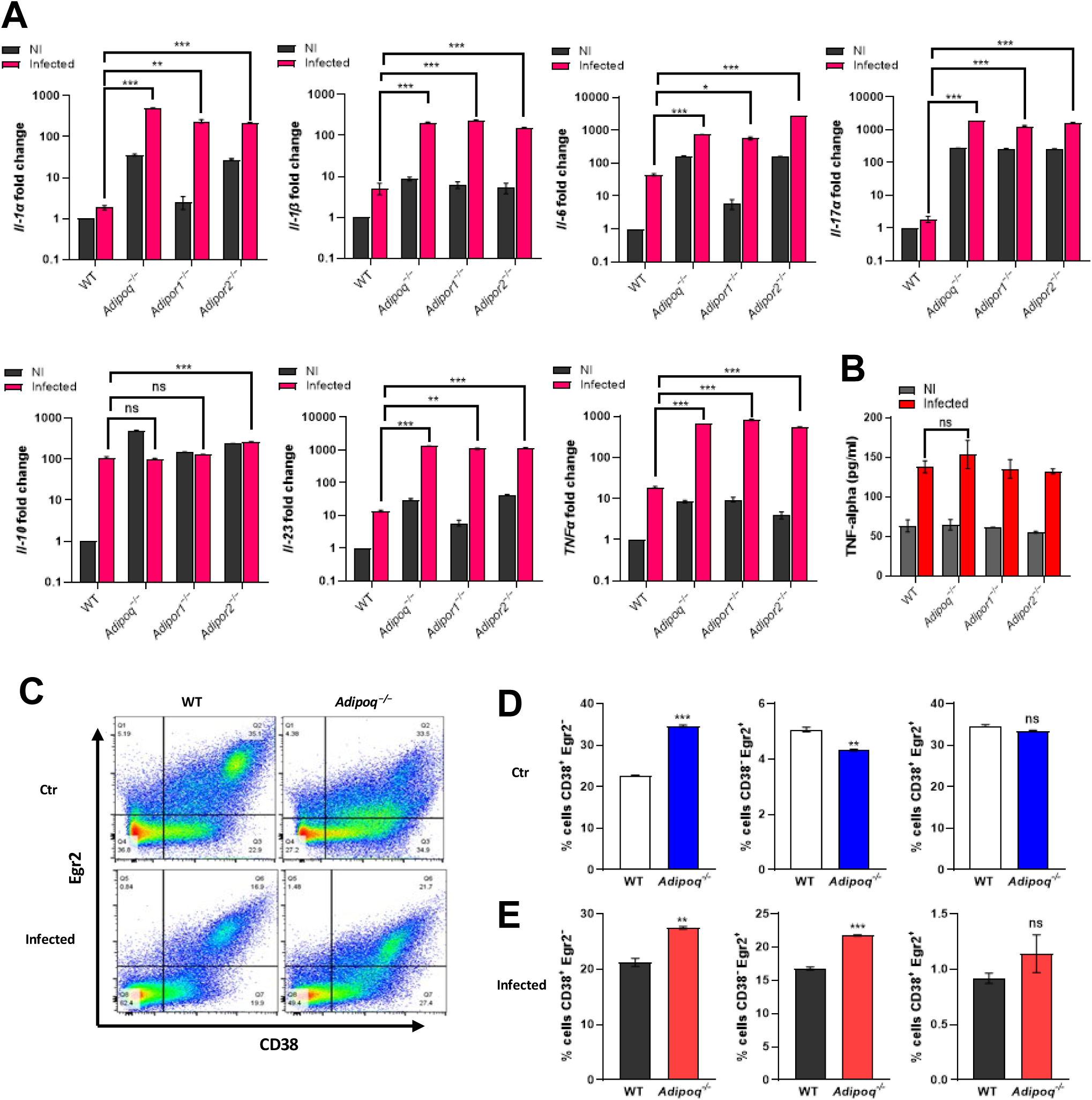
Alveolar macrophages from APN pathway-deficient mice exhibit an inflammatory phenotype. BALF cells from wild-type (C57BL/6), APN-/- (*Adipoq*^−/−^), *AdipoR1*^−/−^, and *AdipoR2*^−/−^mice were collected in PBS/EDTA. For alveolar macrophage differentiation, total BALF cells were plated and cultured with GM-CSF. Alveolar macrophages were challenged with conidia in the ratio 1:9 (AM: conidia) for 10 hours duration as per the details mentioned in Materials and Methods. The data represented is AMs from n=3 mice/group. A. qRT-PCR analysis for mRNA expression of the indicated cytokines. B. TNFα secretion quantified at the protein level by ELISA using *ex vivo* cultured alveolar macrophages. C. Flow cytometry staining of surface CD38 and intracellular Egr2 (D-E) Quantification of (C), showing the proportion of M1, M2 and CD38^+^Egr2^+^ macrophages with CD38^+^Egr2^−^ (putative M1) or CD38^−^Egr2^+^ (putative M2) phenotype in control group (D) and infection group (E). Data are a summary of three independently performed experiments. \**p* < 0.05, \*\**p* < 0.01, \*\*\**p* < 0.001.

### *A. fumigatus* uptake and killing is inhibited in APN-/- AMs

Although APN pathway-deficient mice displayed increased IA severity, fungal burden, and an increased AM inflammatory phenotype, the mechanism of decreased fungal clearance remains unknown. To determine if APN-deficient AMs were deficient in fungal uptake and killing, we used dual viability/tracking fluorescent *Aspergillus* reporter FLARE conidia [15]. to compare changes in conidia uptake and viability in wild-type and APN-/- AMs ex vivo and in vivo. In APN-/- AMs cultured ex vivo, uptake and killing of dual-labeled FLARE conidia was decreased after 6 hours when observed by microscopy (**Fig. 3A** and **Supplemental Video SV1**). When loss of *A. fumigatus* viability was compared by image analysis, killing of FLARE conidia by APN-/- AMs was markedly decreased compared to wild-type AMs (**Fig. 3B**). Uptake and killing were also reduced in APN-/- AMs when analyzed by flow cytometry, even after filtration to remove hyphae (**Figs 3C and 3D**). Reduced killing by APN-/- AMs was confirmed in vivo in FLARE-infected neutropenic mice, with a significant difference in conidial viability by day 3 post-infection (**Figs. 3E and 3F**), while uptake of 1 µm inert particles was not significantly different between wild-type and APN-/- AMs (**Fig. 3G**). Collectively, these results demonstrate that AM fungal uptake and killing are decreased in the absence of APN.

**Fig. 3.**
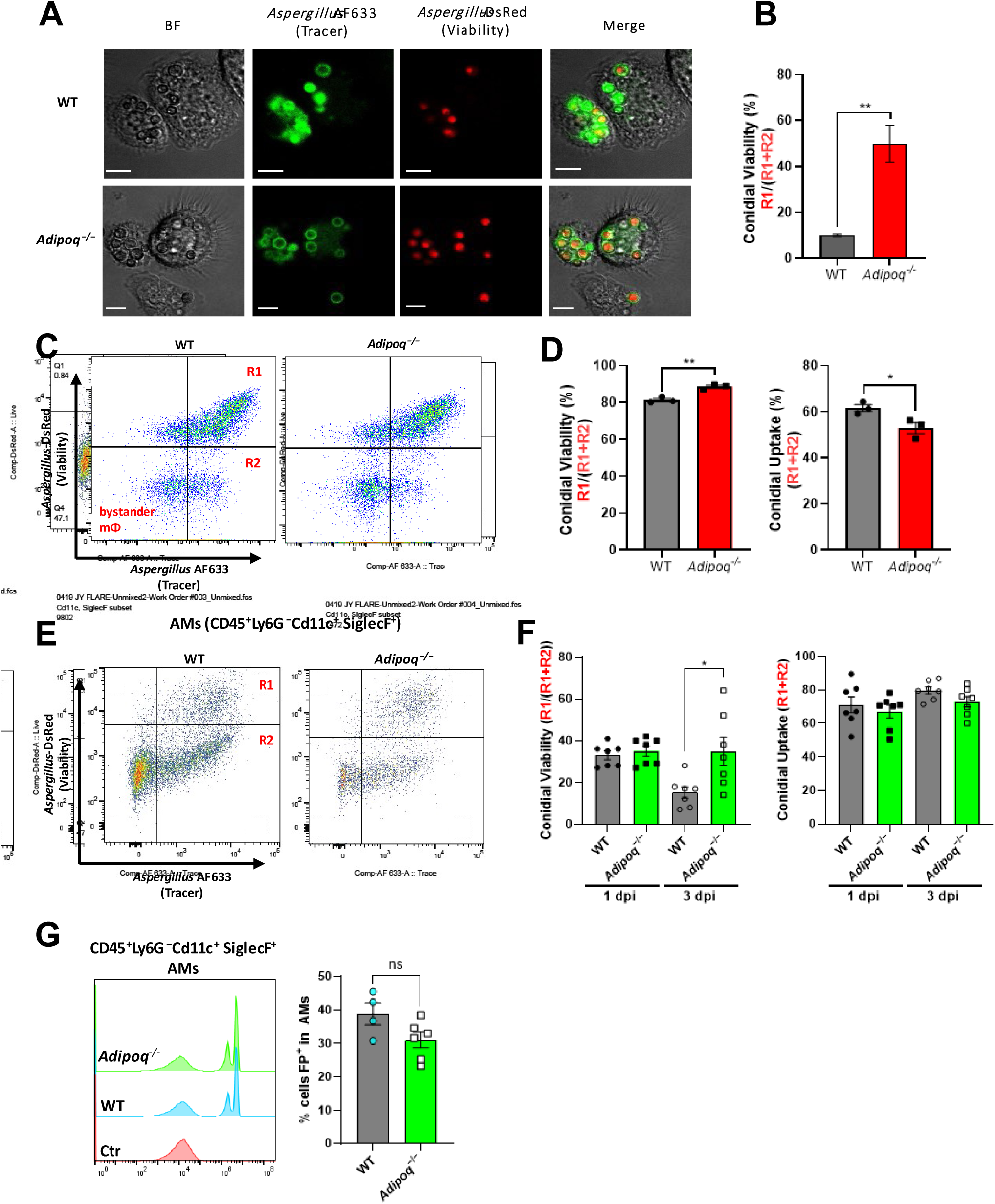
*A. fumigatus* conidial uptake and killing are inhibited in adiponectin-deficient AMs. A. Microscopic analysis of uptake and killing of FLARE conidia with magnification 40x is depicted in Brightfield, Cy5, DsRed channels followed by the merged image. Scale bar is equal to 12µm. B. Quantification of *Aspergillus* viability using microscopy images via calculation of co-localization in imageJ. C. Representative plots that display RFP and AF633 fluorescence intensity of *ex vivo* cultured alveolar macrophages where R1 represents live conidia inside the alveolar macrophages and R2 represents dead/killed conidia. D. Quantification of (C), showing *Aspergillus* viability (R1/(R1 + R2)) and uptake (R1 + R2) in *ex vivo* cultured alveolar macrophages. E. Representative plots that display RFP and AF633 fluorescence intensity of AMs (CD45^+^Ly6G^−^CD11c^+^SiglecF^+^) *in vivo.* Mice of both groups were made immunocompromised and infected with 1-1.5 × 10^7^ conidia. BAL cells were harvested 1 day and 3 days later. F. Quantification of (E), showing *Aspergillus* viability (R1/(R1 + R2)) and uptake (R1 + R2) in alveolar macrophages in neutropenic mice at d1 and d3 p.i. G. Fluorescent particle uptake by alveolar macrophages in vivo. For flow cytometric analysis, neutrophil-depleted mice were challenged with 1.5 × 10^7^ of flow particles. Percentage of flow particle positive (FP+) cells was assessed in AMs (CD45^+^Ly6G^−^CD11c^+^SiglecF^+^) from the wild-type and *Adipoq* ^−/−^ mice by flow cytometry. A-D, data are representative of three independent experiments. E-G, data are a summary of 2-3 independent experiments. \**p* < 0.05, and \*\**p* < 0.01.

### The AdipoR agonist AdipoRon decreases inflammation and fungal burden in APN-deficient mice with IA

A small molecule agonist (AdipoRon) that binds and activates signaling through both AdipoR1 and AdipoR2 improved insulin resistance and glucose intolerance and increased the lifespan of obese mice [16]. A role for AdipoRon in dampening inflammatory responses in human lung macrophages via inhibition of production of TNF, IL-6, and inflammatory chemokine production has also been reported [17]. However, the ability of AdipoRon therapy to improve IA outcomes remained unknown. We treated infected neutropenic wild-type or APN-/- mice with AdipoRon by involuntary aspiration daily for up to eight days (**Fig. 4A**). Although AdipoRon treatment did not significantly improve survival (**Fig. 4B**), lung hyphal growth and fungal burden were reduced in APN-/- mice (**Figs. 4C-4E**). Airway eosinophil recruitment was reduced in AdipoRon treated APN-/- mice, while AMs were increased in both AdipoRon-treated WT and APN-/- mice (**Fig. 4F**). Transcription of IL-1a, IL-6, IL-10, IL-17A, and TNF were also decreased with AdipoRon treatment in APN-/- lungs and decreases in IL-1a and IL-10 were also evident in wild-type mice (**Fig. 4G**). In contrast, IL-12b transcription appeared to be increased by AdipoRon, but only in wild-type lungs. An AdipoRon-mediated decrease in TNF was also confirmed in APN-/- BALF at the protein level (**Fig. 4H**). These results indicate that APN pathway stimulation by AdipoRon decreases lung fungal burden and dampens lung inflammation in APN-/- mice with IA.

**Figure 4.**
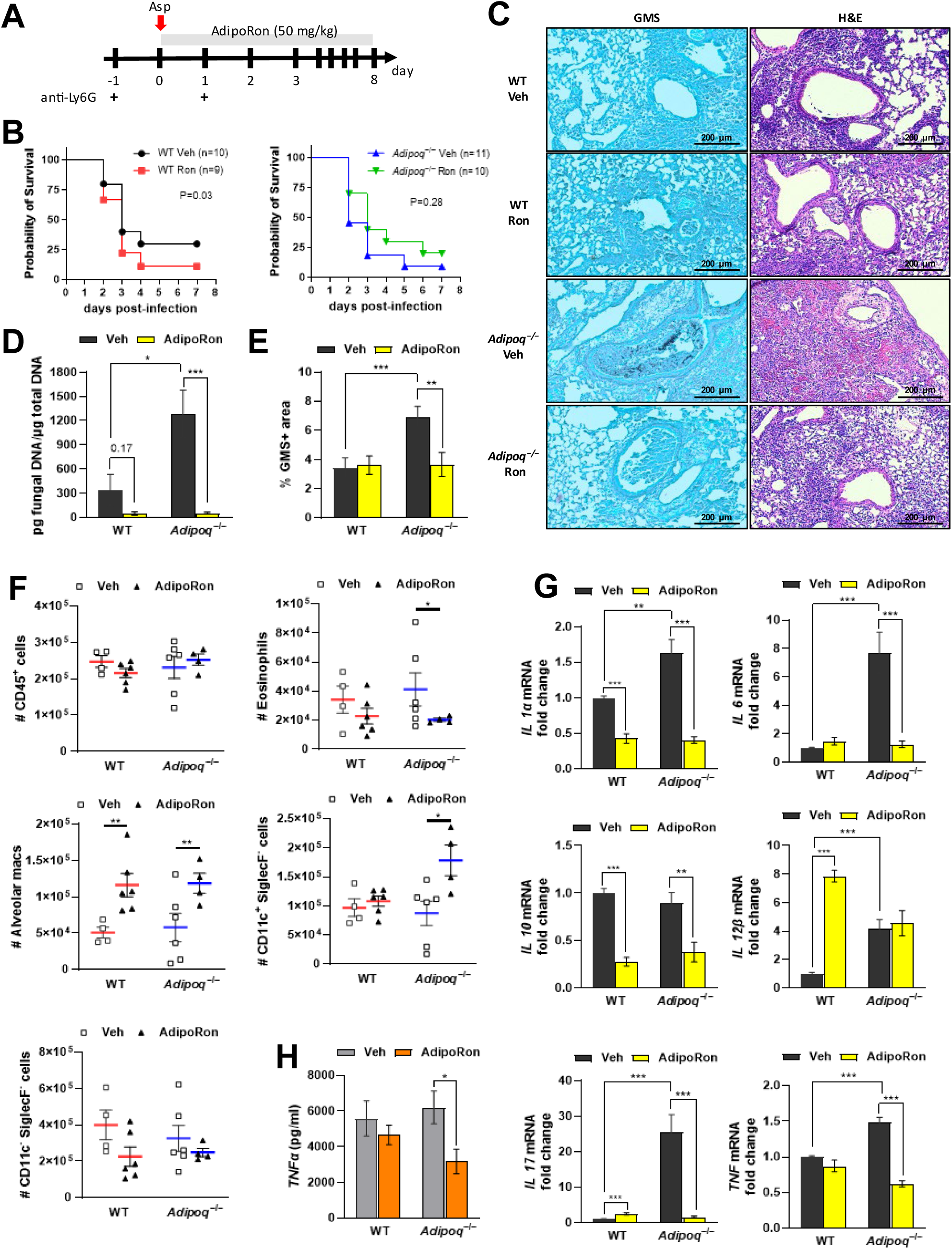
The AdipoR agonist AdipoRon improves decreases inflammation and fungal burden in APN-deficient mice with IA. Wild-type (C57BL/6), and *Adipoq*^−/−^ mice were neutrophil depleted and involuntarily aspirated *A. fumigatus* conidia as described in Materials and Methods. A. Experimental time course for neutropenic mouse model of infection and AdipoRon treatment. B. Effect of AdipoRon on the survival rate of WT and *Adipoq*^−/−^ mice. *N* = 9-11 mice per group. C. C. Representative GMS and H&E lung sections. D. Fungal burden determined by quantitative PCR of fungal DNA from lung homogenates. E. Fungal burden determined by quantification of GMS staining. F. Total number of CD45^+^ cells, eosinophils (CD45^+^Ly6G^−^CD11c^−^SiglecF^+^), AMs (CD45^+^Ly6G^−^CD11c^+^SiglecF^+^), CD11c^+^SiglecF^−^ (CD45^+^Ly6G^−^ CD11c^+^SiglecF^−^), and CD11c^−^SiglecF^−^ (CD45^+^Ly6G^−^CD11c^−^SiglecF^−^) cells isolated from the mice with IA as determined by flow cytometry. *N* = 4-6 mice per group. G. qRT-PCR analysis for mRNA expression of the indicated cytokines. H. TNFα secretion in BALF quantified at the protein level by ELISA. Data are a summary of 2-3 independently performed experiments. \**p* < 0.05, \*\**p* < 0.01, \*\*\**p* < 0.001.

### AdipoRon increases fungal killing and inhibits the inflammatory phenotype of APN-/- AMs

Although AdipoRon treatment influenced lung fungal growth and dampened inflammation in APN-/- mice with IA, the specific effect of AMs remained unknown. To determine the effect of AdipoRon on AM phenotype and fungal uptake and killing, we infected WT or APN pathway-deficient AMs with Af293 or FLARE conidia after 24 hours treatment with AdipoRon or control vehicle (containing DMSO), and harvested AMs ten hours later for microscopic, flow cytometric, qRT-PCR, and ELISA analyses (**Fig. 5A**). When quantified by microscopy, conidia viability was decreased by AdipoRon treatment of APN-/- but not wild-type AMs (**Fig. 5B**). Conidia viability was also reduced in APN pathway-deficient AMs by AdipoRon when measured by flow cytometry after cell straining to remove hyphae (**Fig. 5C**). Inflammatory cytokine transcription of IL-1α, IL-1β, and IL-6 were all decreased by AdipoRon treatment in APN-/- AMs, while the anti-inflammatory cytokine IL-10 was not affected by AdipoRon regardless of APN sufficiency (**Fig. 5D**). Secretion of TNF protein was also significantly decreased in AdipoRon-treated APN-/- AMs (**Fig. 5E**). These results indicate that AdipoRon treatment acts on APN-/- AMs to improve fungal killing and inhibit an inflammatory phenotype.

**Fig. 5.**
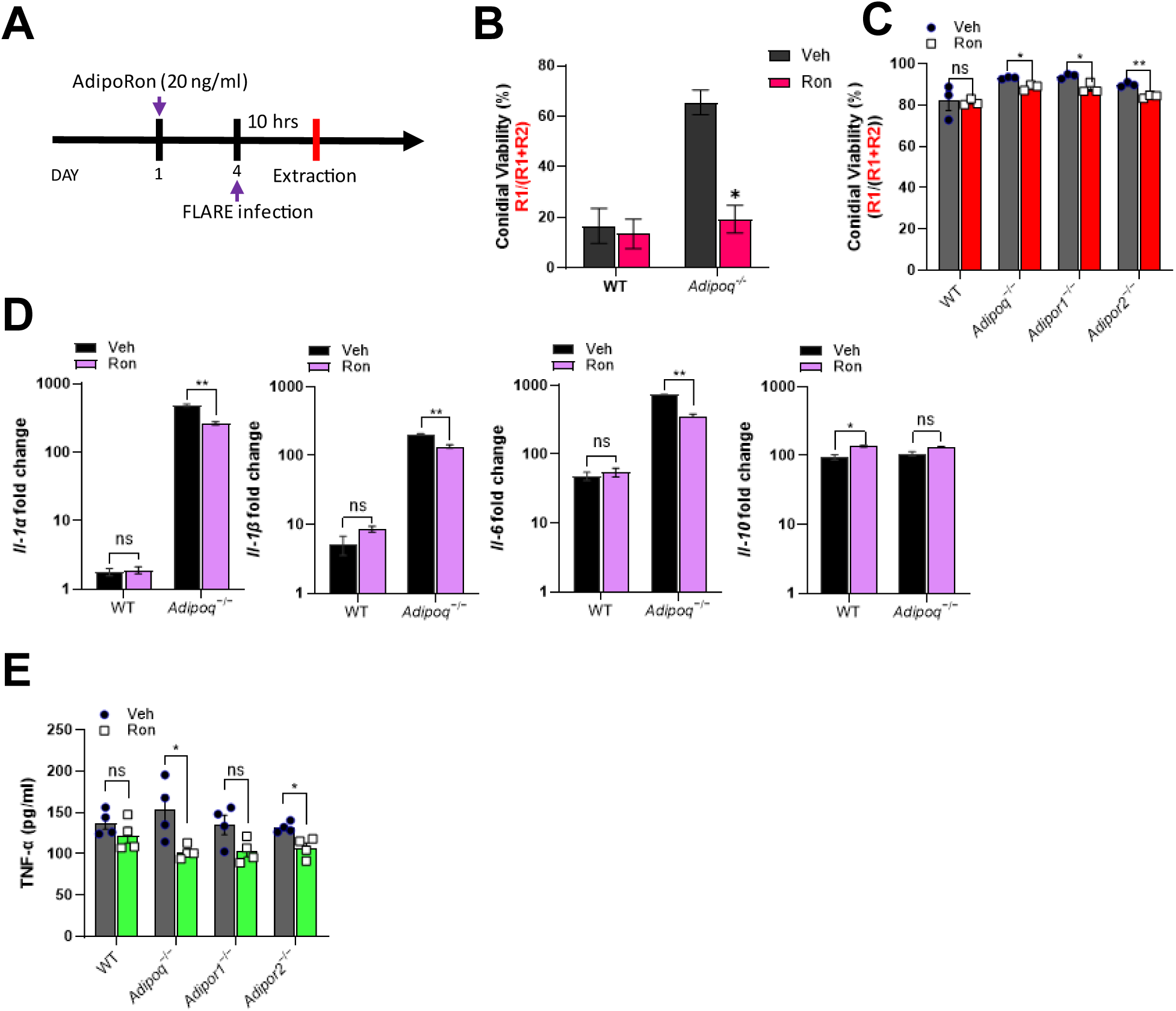
AdipoRon increases AdipoR-dependent AM fungal killing and reverses inflammatory phenotype. A. Experimental time course for *ex vivo* cultured AMs challenged with conidia with or without AdipoRon treatment. B. Microscopic image quantification of *Aspergillus* viability (R1/(R1 + R2)) in *ex vivo* cultured AMs by using FLARE conidia. The infection was done ex-vivo in 1:1 ratio (AMs: FLARE) for 15 hours. C. Flow cytometric quantification of *Aspergillus* viability. D. qRT-PCR analysis for mRNA expression of the indicated cytokines. E. TNF secretion quantified at the protein level using *ex vivo* cultured AMs by ELISA. Data are representative of three independently performed experiments. \**p* < 0.05, \*\**p* < 0.01, \*\*\**p* < 0.001.

### AdipoRon-enhanced killing of *A. fumigatus* requires both AdipoR1 and AdipoR2

Since AdipoRon enhances AM-mediated fungal killing, we aimed to confirm the requirement of AdipoR1 and AdipoR2 by using siRNA to target expression of AdipoR1 in wild-type, APN-/-, and AdipoR2-/- AMs (**Fig. 6A**). Decreased AM transcription of *Adipor1* and *Adipor2* by infection was partially restored by AdipoRon treatment (**Fig. S2E**). Knockdown of AdipoR1 was confirmed by flow cytometry to reduce expression to identical levels observed in AMs from AdipoR1-/- mice (**Figs. 6B and 6C**). Moreover, fungal killing was improved by AdipoRon treatment in wild-type, APN-/-, AdipoR1-/- (KO mice and siRNA), and AdipoR2-/- AMs, but was not improved with siRNA-targeted depletion of AdipoR1 in AdipoR2-/- AMs (**Fig. 6D**). These results show that both canonical APN receptors are critical for AdipoRon-mediated enhancement of AM fungal killing.

**Fig. 6.**
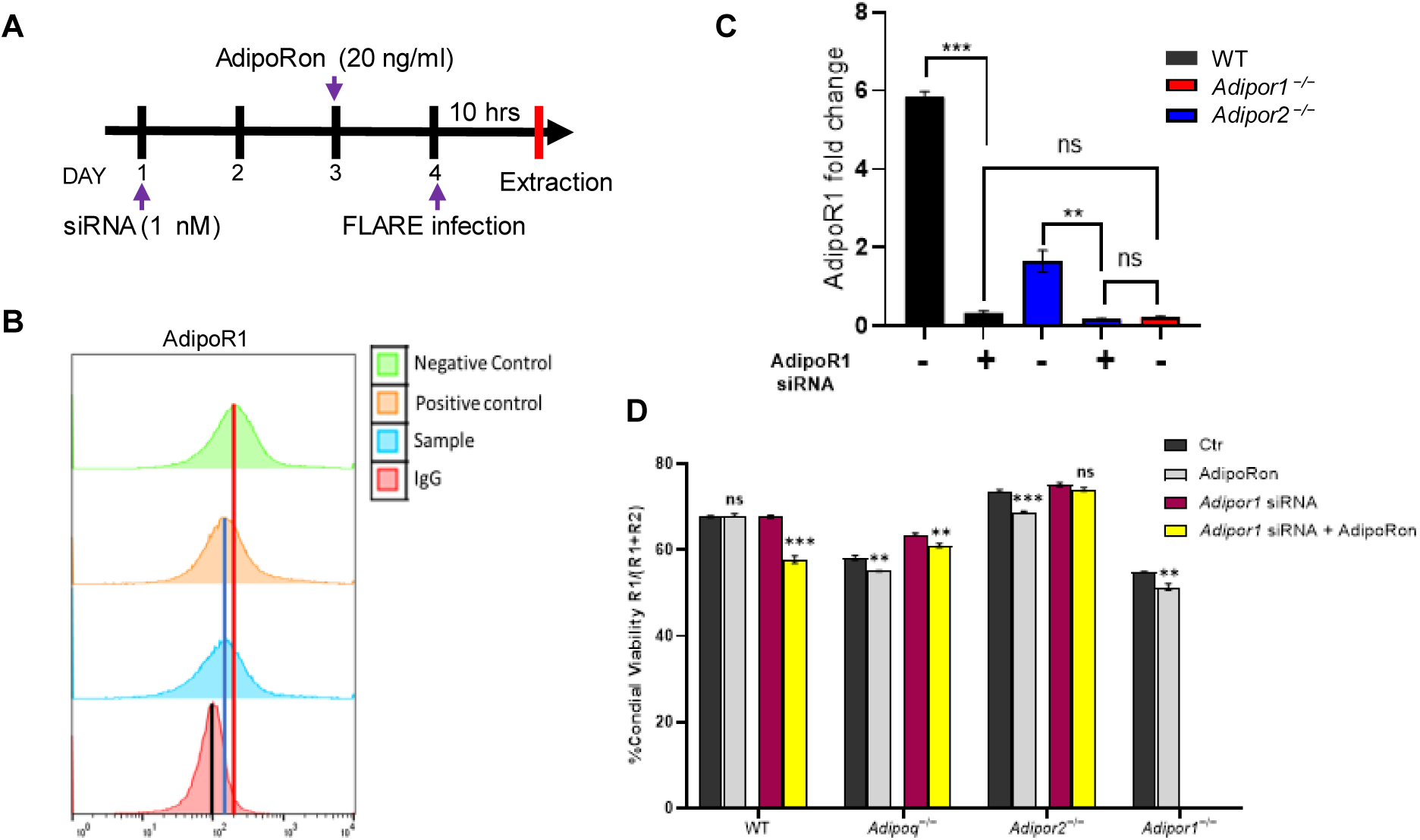
AdipoRon-enhanced killing of *A. fumigatus* requires both AdipoR1 and AdipoR2. A. Schematic timeline course for primary AM cultures treated with *AdipoR1* siRNA, details in Methods section. B. Histogram graphs obtained from flow cytometry analysis. The positive control represents the ideal *Adipor1* knockdown control, while the negative control is in the absence of functional SiRNA. C. qRT-PCR analysis for mRNA expression of the *adipor1* in AMs from the APN-deficient mice with siRNA treatment. D. Conidial viability (R1/(R1 + R2)) in *ex vivo* cultured AMs with siRNA and/or AdipoRon by flow cytometry Data are representative of two independently performed experiments. \*\**p* < 0.01 and \*\*\**p* < 0.001.

### Adiponectin promotes LC3-associated phagocytosis of *A. fumigatus* conidia

To confirm the AdipoRon-mediated phenotypic shift in AMs, we compared AdipoRon treatment with vehicle in infected and uninfected APN-/- AMs by global RNAseq analysis. In uninfected AMs, more genes were upregulated in response to AdipoRon treatment, while the opposite was observed in AMs after 12 hours post-infection, with AdipoRon treatment associated with gene silencing (**Fig S3A**). Inflammatory/M1-associated gene expression was suppressed by AdipoRon treatment in infected AMs, including *Il1a*, *Il1b*, *Ccl2*, *Cd38*, *Tlr4*, and *Hif1a*, while the anti-inflammatory *Tgfb* was increased, as were autophagy genes, including *Map1lc3a* and *Map1lc3b* genes that are critical for LAP (**Fig. S3B**). In contrast, inflammatory gene expression appeared to be increased by AdipoRon in uninfected AMs. GSEA pathway analysis confirmed inhibition of cytokine signaling by AdipoRon in infected AMs and also identified activation of gene pathways associated with oxidative phosphorylation (**Fig. S3C**), a metabolic pathway favored by M2 macrophages [18]. Our RNAseq results thus confirm an AdipoRon-mediated inhibition of the inflammatory/M1 phenotype of APN-/- AMs in response to fungal infection, and identify a potential role for autophagy induction.

Since LAP is an important mechanism of killing of *A. fumigatus* by macrophages [19], we aimed to determine if phagosomal LC3 activation is increased in APN-/- AMs by AdipoRon treatment. Using immunofluorescence microscopy staining of LC3, we observed clear colocalization of LC3 staining and DS-RED+(viable) conidia in AdipoRon-treated APN-/- AMs (**Fig. 7A**), with visibly less colocalization in untreated AMs that was confirmed by image quantification (**Fig. 7B**). The median fluorescence intensity (MFI) of LC3+ macrophage staining was also increased by AdipoRon in both wild-type and APN-/- AMs as measured by flow cytometry (**Fig. 7C**). Collectively, these results demonstrate a role for adiponectin in LAP in AMs in response to *A. fumigatus* conidia.

**Fig. 7.**
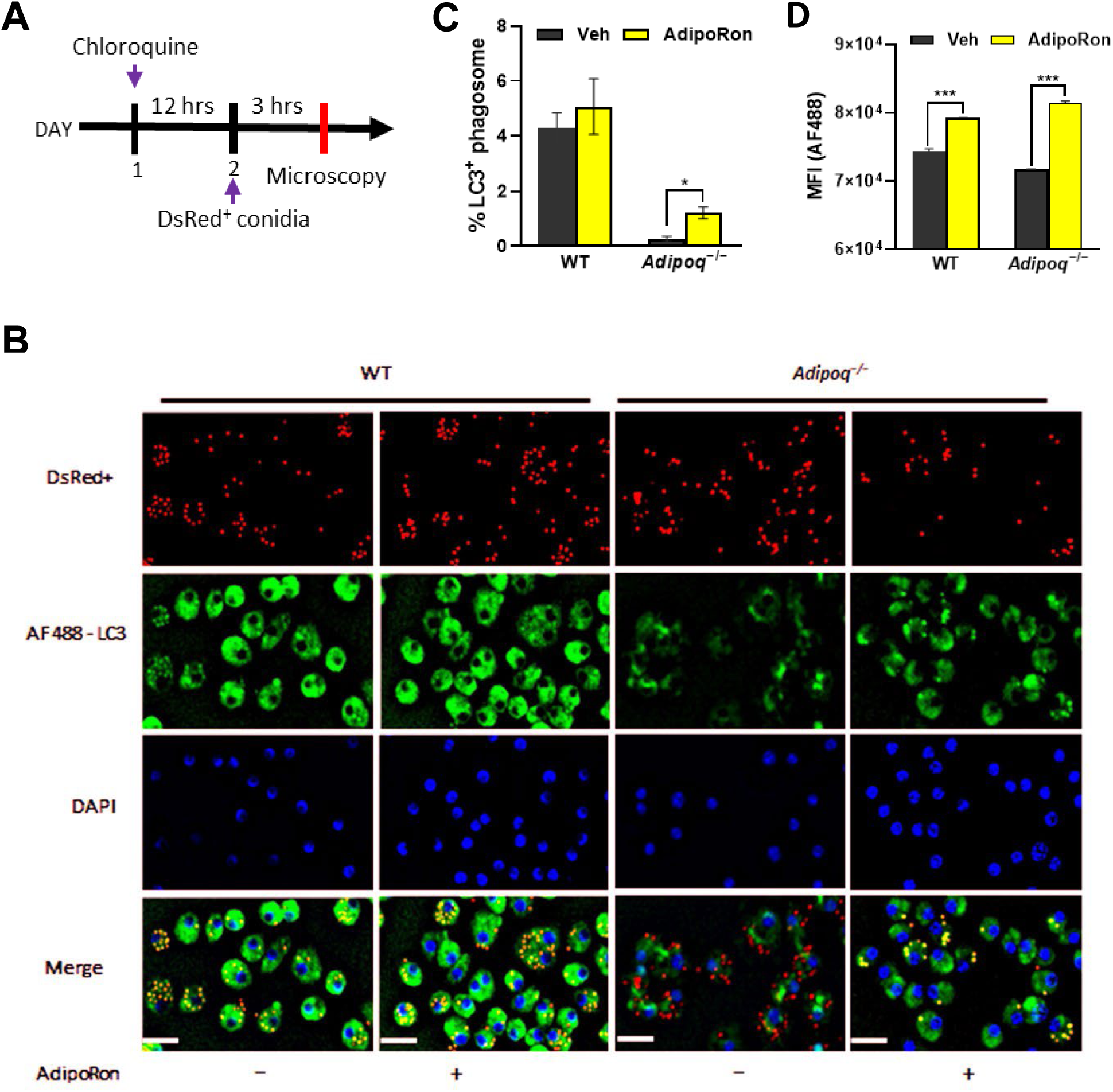
Adiponectin promotes LC3-associated phagocytosis of *A. fumigatus* conidia. A. Schematic timeline course for LC3-associated phagocytosis analysis. B. Microscopy of LAP-dsRed conidia and LC3^+^ AMs at a magnification of 20X in *ex vivo* cultured AMs from infected WT and *Adipoq*^−/−^ mice with or without AdipoRon. The images from DsRed, FITC, DAPI channels are presented followed by the merged image. Scale bar: 35µm. C. Microscopic quantification of LC3^+^ phagosome (%) in *ex vivo* cultured AMs from infected WT and *Adipoq*^−/−^ mice with or without AdipoRon. D. The mean fluorescent intensity (MFI) obtained by flow cytometry indicates the expression level of LC3-AF488 in *ex vivo* cultured AMs from infected WT and *Adipoq*^−/−^ mice with or without AdipoRon. Data are representative of two independently performed experiments. \**p* < 0.05 and \*\*\**p* < 0.001.

## Discussion

To our knowledge, our study is the first to identify a role for adiponectin in LC3-associated phagocytosis (LAP) in response to infection. Previous studies reported roles for adiponectin in LC3-mediated autophagy in skeletal muscle cells [20,21], cardiomyocytes [22], or macrophages in a mouse model of atherosclerosis [23]. In addition, AM autophagy is critical for prevention of spontaneous lung inflammation driven by airway microflora [24]. Although others have identified a role for the LC3 pathway in efferocytosis by macrophages [25], and although adiponectin promotes macrophage efferocytosis [26,27], a direct connection between adiponectin, LC3/autophagy and macrophage efferocytosis is lacking. Since macrophage autophagy and removal of apoptotic cells are important mechanisms to limit lung inflammatory pathology [24,27,28], it will be of interest in future studies to determine their role in adiponectin-mediated dampening of inflammation in IA.

A number of studies have defined a critical role for LAP in protection from *A. fumigatus* infection [19,29–31]. LAP dampens inflammation in response to *A. fumigatus* by inhibition of the NLRP3 inflammasome and IL-1b via activation of DAPK1 [29]. In our study, we observed that AdipoRon decreased transcription of IL-1α and IL-1β genes in APN-/- AMs. Interestingly, blockade of IL-1 receptor signaling (by IL-1ra) restored autophagy/LC3 recruitment in NADPH oxidase-deficient macrophages, a hallmark myeloid cell deficiency of Chronic Granulomatous Disease [32]. Since both AdipoRon and IL-1ra promote LC3 activation, it will be important in future studies to determine if AdipoRon could restore autophagy in NADPH oxidase-deficient macrophages, and to identify shared regulatory pathways.

A previous study reported that LAP-associated fungal clearance was inhibited by fungal melanin, a virulence factor of *A. fumigatus* [30]. How APN stimulates LAP in response to *A. fumigatus* and the ability to overcome LAP interference by fungal melanin remain unknown. Furthermore, we observed that chitin inhalation resulted in decreased production of lung adiponectin and AdipoR1 expression in murine airway cells [33]. It will be of interest to determine the effect of AdipoRon on recognition of multiple fungal pathogen-associated molecular patterns by WT and APN pathway-deficient AMs, specifically in the context of LC3 activation.

Adiponectin has two canonical signaling receptors with broad, yet different cell/tissue distribution [34]. We observed that both AdipoR1 and AdipoR2 contribute to adiponectin-mediated protection from IA and dampening of inflammatory pathology. However, some differences were evident, as AdipoR1-/- mice with IA exhibited the highest levels of lung inflammatory cytokine transcription that was not as clear in AdipoR1-/- AMs, suggesting that immune pathology in AdipoR1-/- mice was not driven solely by the lack of AdipoR1 signaling in AMs, but also by other immune and resident lung cells, with a contribution of potentially enhanced signaling through AdipoR2. AMs express both AdipoRs, and their expression was modulated by *A fumigatus* infection, most significantly for AdipoR1, which was increased in response to infection in WT AMs but decreased in APN-/- AMs. Furthermore, both AdipoR1 and AdipoR2 were required for AdipoRon-mediated improvement in *A. fumigatus* killing, as AdipoRon was not effective in AdipoR1/R2-/- AMs, confirming a role for both receptors in APN-mediated enhancement of AM function. However, the relative contribution of AdipoR-expressing AMs in adiponectin-mediated protection from infection remains unclear, and thus remains a subject of future investigation.

AMs are dynamic cells, and their activation and polarization falls on a spectrum from M1 to M2, as these are designations and not indicative of a static phenotype [35,36]. In our study, AM treatment with AdipoRon decreased expression of M1 genes and activated M2 genes in *A. fumigatus*-infected APN-/- AMs, although these phenotypes are not identical to previously published studies using classical activation stimuli, such as LPS/IFNg for M1 and IL-4 for M2 [37,38]. It is thus more appropriate to refer to APN-/- AMs as ‘M1-like’ and AdipoRon-treated AMs as ‘M2-like’ cells [39,40]. Perhaps more importantly, AdipoRon treatment of APN-/- AMs resulted in activation of genes associated with oxidative phosphorylation that is associated with M2-like macrophages, in contrast to glycolysis-favoring M1 macrophages [18]. In support of increased oxidative metabolism for AdipoRon-treated APN-/- AMs, arginase-2 (*Arg2*) expression was also increased, and mitochondrial *Arg2* was recently identified as critical for induction of oxidative phosphorylation in inflammatory macrophages [41]. Collectively, the results of our study confirm a role for APN in skewing AMs toward an M2-like phenotype.

Obesity is widely accepted as an inflammatory disease characterized by adipose tissue inflammation and decreased APN and APN receptor expression [6,42–45], yet the effect of obesity-related APN pathway-inhibition in response to *A. fumigatus* infection is not known. In response to infection with *Listeria monocytogenes*, both obese and APN-/- mice exhibited attenuated clearance that was characterized by an inflammatory phenotype in bone marrow macrophages that was partially restored by AdipoRon [7]. Furthermore, obesity is associated with an increased inflammatory phenotype in alveolar macrophages [46]. It is therefore possible that stimulation of the APN pathway with AdipoRon will improve AM killing of *A. fumigatus* in obese mice, an outcome that would support the APN pathway as a potential therapeutic route in IA patients with low APN levels.

## Acknowledgements

We would like to thank Azriel Manalaysay, Alsace Key, and Gwen Schmiedel for assistance with animal care and genetic screening, Brielle Batch for technical assistance, Takuya Akiyama for assistance with confocal microscopy, Americo Lopez-Yglesias for assistance with flow cytometry, and Tobias Hohl for review of this manuscript. The study was supported by National Institutes of Health NIH R21AI163574 (ST) and T32 AI060519 (NBA; IUSM Department of Microbiology and Immunology, American Lung Association Innovation Award #IA-690880 (ST), and the Indiana Clinical and Translational Sciences Institute, funded in part by grant # UM1TR004402 from the National Institutes of Health, National Center for Advancing Translational Sciences, Clinical and Translational Science Award (ST).

## Author contributions

Conceptualization, S.P.T., H.G., J.L.; Methodology, J.L. and H.G.; Investigation. H.G., J.L., and N.B.A.; Resource; J.L., N.B.A., and H.G.; Writing - S.PT, J.L and H.G.; Visualization, H.G, J.L.; Funding Acquisition. S.P.T.

## KEY RESOURCES TABLE

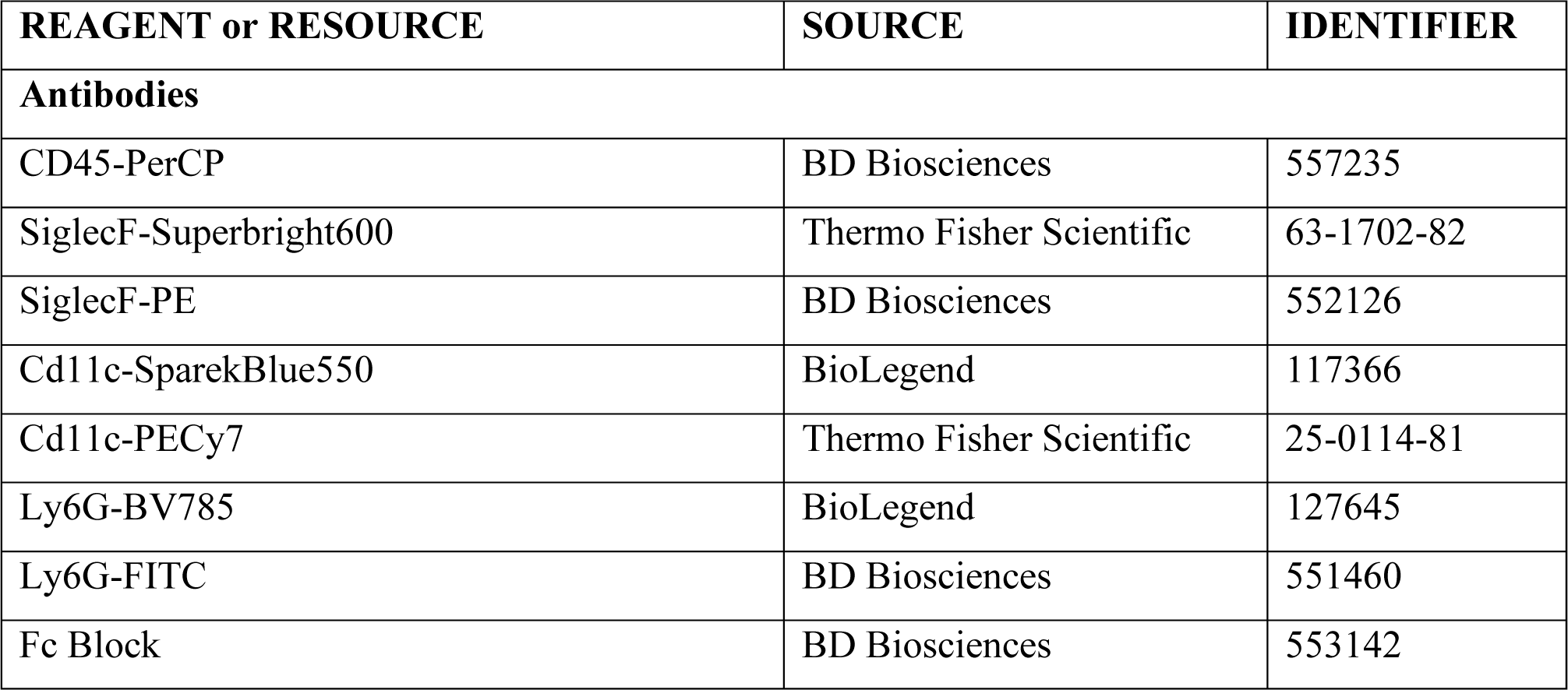
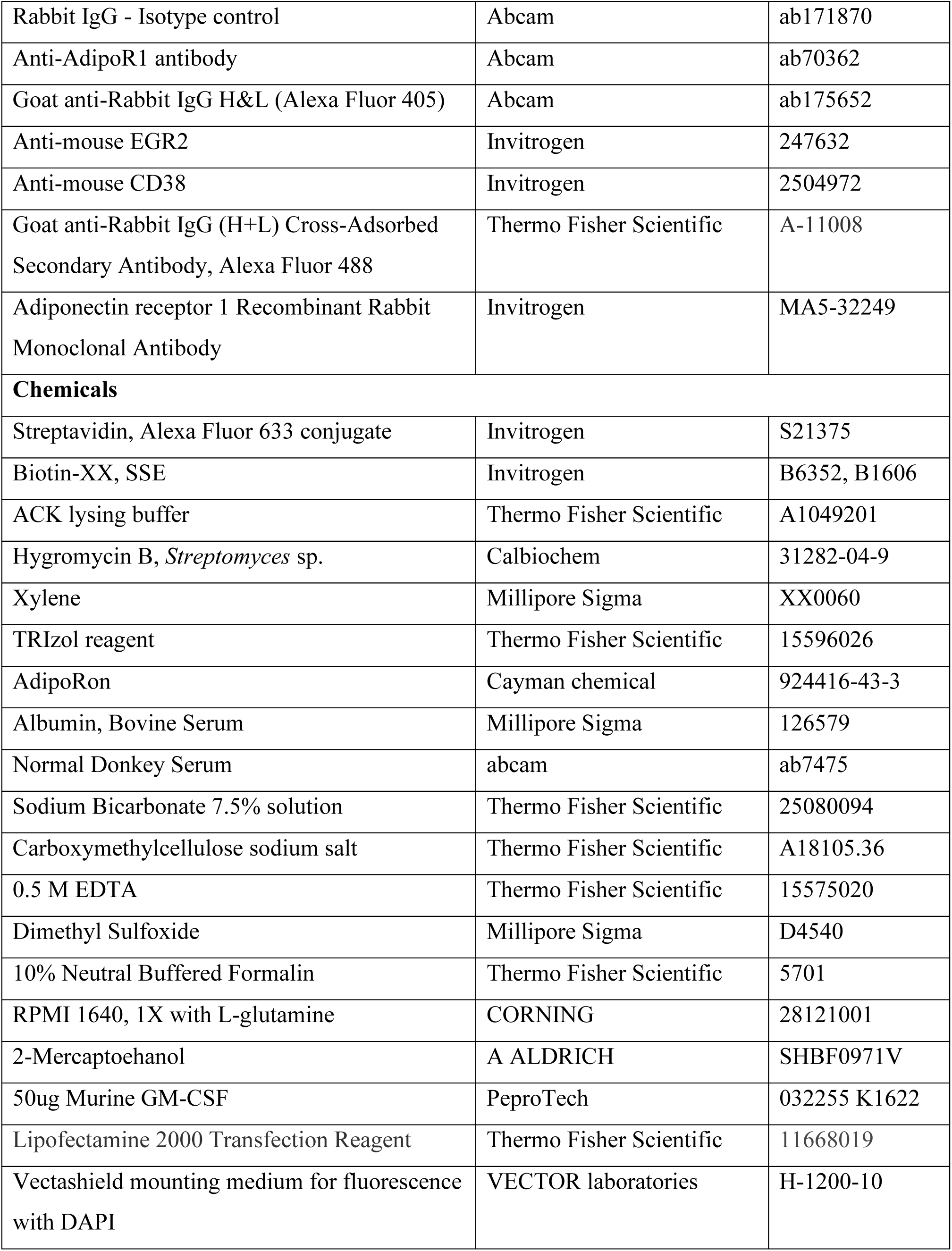
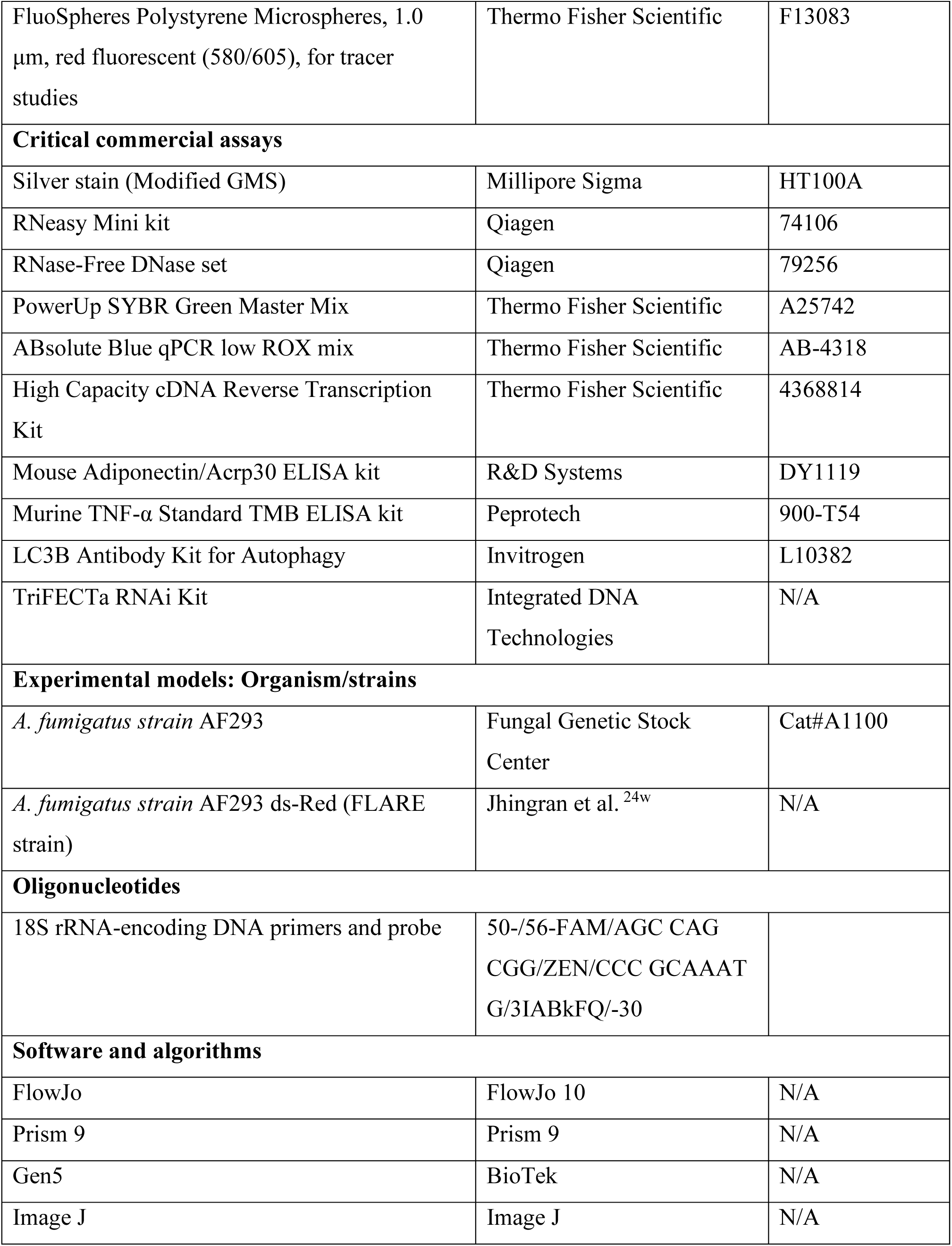

## RESOURCE AVAILABILITY

### Lead contact

Further information and requests for resources and reagents should be directed to and will be fulfilled by the lead contact, Steven P. Templeton.

### Material Availability

All materials within the paper are available from the corresponding author upon reasonable request. This study did not generate new reagents.

### Data and Code Availity

The RNAseq data are available at Gene Expression Omnibus (Accession #GSE268782).

## Materials and Methods

### Mice

C57BL/6 and adiponectin-deficient (*Adipoq*−/−) and *AdipoR2*+/− mice were obtained from The Jackson Laboratory, while AdipoR1+/- mice were obtained from the Mutant Mouse Resource and Research Center (MMRRC). To obtain homozygous deletion mice for *AdipoR1*−/−, and *AdipoR2*−/− mice, heterozygous mice were breed with F1 animals heterozygous and offspring were screened by PCR, with loss of protein confirmed in a subset of animals by flow cytometry. All animal handling and experimental procedures were performed in accordance with the recommendations found in the Guide for the Care and Use of Laboratory Animals of the National Institutes of Health. The work in this study was approved by the Institutional Animal Care and Use Committee of the host campus of Indiana University School of Medicine-Terre Haute, Indiana State University.

### Fungal strains and cultivation

The clinical isolates AF293 was previously obtained from the Fungal Genetics Stock Center and grown on Malt Extract Agar plates at 22°C. To generate FLARE conidia, AF293-dsRed conidia (provided by Dr. Tobias Hohl, Memorial Sloan-Kettering) were rotated in 0.5 mg/ml Biotin XX, SSE in 1 ml of 50 mM NaHCO_3_ for 2 h at 4°C, washed with 0.1 M Tris-HCl (pH 8.0), incubated with 0.02 mg/ml Af633-streptavidin for 30 min at RT, and resuspended in PBS for use [47].

### Fungal aspiration and infection

Neutrophils were depleted by i.p. injection of 0.5 mg of α-Ly6G (1A8 clone) 24 h pre- and post-infection for invasive pulmonary aspergillosis mice model, as described [8]. Isoflurane-anesthetized mice involuntarily aspirated 50 μl of suspension including 1 – 2 × 10^7^ of conidia. For survival test, infected mice were monitored for 8 days post infection. For further analyses, a subset of infected mice were euthanized with sodium pentobarbital and mouse BAL cells and lungs were harvested 24 h or 72 h later for further analyses.

### Histology

At 3 days post infection, lungs were harvested after perfusion with 5 ml of PBS followed with 5 ml of 10 % phosphate buffered formalin and inflated with formalin. Lung tissue was embedded in parafilm after dehydration and sectioned in 4 μm slices using xylene and ethanol and stained with modified Gomori modified methanamine silver (GMS) and hematoxylin and eosin (H&E) staining [8]. Quantification of GMS staining to determine fungal burden was performed as previously described [48].

### Quantitative PCR fungal burden assay

At 72 h post infection, lungs were harvested, frozen in liquid nitrogen and lyophilized. Genomic DNA was extracted from homogenized lung tissue with DNA extraction buffer for *Aspergillus* nucleic acids and subsequent phenol/chloroform extraction. Quantitative PCR (qPCR) fungal burden assay was performed using 0.2 μg of genomic DNA with 18S rRNA-encoding DNA primers and probe sets with a modified probe quencher (50-/56-FAM/AGC CAG CGG/ZEN/CCC GCAAAT G/3IABkFQ/-30) [49]. qPCR was performed using the CFX Connect Real-Time System. Threshold values were used to calculate the corresponding fungal DNA content in the lung tissues.

### Tissue RNA extraction and gene expression analysis

Total RNA was extracted from homogenized lungs in TRIzol reagent. Following the upper phase, further RNA purification was performed using a Qiagen RNeasy column and DNase treatment per the manufacturer’s recommendations [50]. Mouse β-actin was used as the housekeeping gene for cytokine analysis in the murine model. Most primers were designed using the Primer3 software (version 0.4.0) from the whole sequence available in GenBank.

### BALF Flow cytometric analysis

BALF cell composition was determined by flow cytometric analysis with staining of surface markers. In brief, BALF was centrifuged and the cell pellet was washed with 1 ml of FACS buffer (PBS, 5% FBS, 50 mM EDTA). The washed pellet was lysed of red blood cells with ACK lysis buffer and stained with 10% rat serum, Fc receptor blocking Ab, and antibodies: rat anti-mouse Ly6G-FITC or -BV785, rat anti-mouse SiglecF-PE or -Superbright600, rat anti-mouse CD45-PerCP, and rate anti-mouse CD11c-PE-Cy7 or -SparkBlue550 [50,51]. For staining of AdipoR1, the washed pellet was resuspended in blocking solution containing 10% donkey serum, Fc receptor blocking Ab, and 1% Bovine serum albumin in PBS for 30 min, incubated with the primary AdipoR1 antibody for 1 h and incubated with the IgG secondary antibodies [33]. Analyses were performed by flow cytometric analysis on a Guava EasyCyte HT system and Cytek. Flow cytometric data were analyzed using FlowJo software.

### BALF ELISA analysis

Serum and BALF supernatants were collected after centrifugation. ELISA analyses were performed for TNF-α cytokine (PeproTech) and Adiponectin (R&D Systems) as described by the manufacturer’s protocol.

### Alveolar macrophage extraction and culture

Seven to 8 weeks old mice were euthanized, and an 18-gauge catheter was inserted into the trachea and alveolar macrophages were harvested from mouse lungs with sterile filtered PBS containing 2 mM EDTA (diluted 1:250 from 0.5 M EDTA stock solution) and 0.5% Fetal bovine serum (FBS) in 5ml BAL fluid. Cells were separated from the BALF by centrifugation at 400 × *g* for 6 min at 4°C and the cells were suspended in RPMI 1640 supplemented with penicillin (100 U/ml), streptomycin (100 U/ml), 5% heat-inactivated FBS, 20ng/ml GM-CSF and 143mM beta-mercaptoethanol. BALF cells were plated in Corning Costar Flat Bottom 12 well Cell Culture Plates at a concentration of 5 × 10^5^ cells per well. The cells were allowed to adhere for 24 hours at 37°C under a humidified atmosphere with 5% CO2 and were washed with PBS thrice and cell culture media was added, allowing only the adherent AMs to culture in the wells. The cell culture media was changed on day 3, day 6 and day 9 of the cultures. AMs were fully confluent and ready for treatment and infection on day 10. The wells for divided into DMSO control, AdipoRon treatment with AF293/FLARE infection and AF293/FLARE infection groups.

### Quantitative RT-PCR for ex-vivo gene expression analysis

AMs were treated with 20 uM AdipoRon or vehicle-treated for 24 hours, followed by infection with AF293 fixed and swollen conidia using RPMI 1640 medium at a concentration of 9 conidia per cell for RT-PCR. After 10 hours of infection, cells were ready for RNA extraction and cDNA synthesis. RNA extraction from AMs was done using Qiagen RNeasy Mini kit, following the manufacturer’s protocol.

### AM supernatant ELISA

After culturing for 10 days, AMs were grouped into uninfected, vehicle (DMSO), infected and AdipoRon + infected groups. After the infection at a concentration of 9 conidia per cell with fixed and swollen AF293 conidia for 10 hours, the supernatants were collected and analyzed for TNF-alpha expression.

### AM flow cytometric analysis

#### FLARE uptake and killing

Conidia was harvested from 7-10 days old Minimal Media (GMM) plates. AF293-dsRed conidia were rotated in 0.5 mg/ml Biotin XX, SSE in 1 ml of 50 mM NaHCO3 for 2 h at 4°C, washed with 0.1 M Tris-HCl (pH 8.0), incubated with 0.02 mg/ml Af633-streptavidin for 30 min at RT, and resuspended in PBS for use. For in-vitro conidial uptake and killing assays, 5×10^5^ FLARE conidia were added to 5×10^5^ alveolar macrophages (1:1). After 10 hours alveolar macrophages were harvested using cell scraper with slight force from the wells in 500 ul ice cold FACS buffer per well and centrifuged at 400 × *g* for 6 min at 4°C. The cell pellet was washed with 1 ml of FACS buffer (PBS, 5% FBS, 50 mM EDTA) and the samples were passed through 70µm filter to clear hyphae and have clear sample. The cells were analyzed by the flow cytometry for dsRed (Ex/Em: 558nm/583nm) and AF633 (Ex/Em: 631nm/650nm) fluorescence followed by gating using FlowJo.

#### M1/M2 phenotype study

For M1/M2 phenotype analysis cells were stained with 10% rat serum, Fc receptor blocking Ab and antibody: *CD38 Monoclonal Antibody (90), APC-eFluor™ 780, eBioscience™* for 15 min at room temperature. For staining with the antibody: *EGR2 Monoclonal Antibody (erongr2), PE-Cyanine7, eBioscience™* cells were permeabilized using Cytoperm and processed for staining.

### AM Microscopy

#### FLARE uptake and killing

Alveolar macrophages were plated in Cellvis 35 mm Glass bottom dish with 14 mm micro-well and were infected with FLARE/DsRed at a 1:1 ratio. Cells were live imaged after 6 hours on a ZEISS LSM780 confocal microscopy.

### LC3 phagocytosis assay

On day 10, cells were treated with 20 uM AdipoRon for 24 hours followed by treatment with 90µm chloroquine for 13 hours. Cells were then infected with DsRed+ conidia for 120 min and then fixed with 3.7% formaldehyde in PBS for 15 min at room temperature. LC3B antibody kit for autophagy was used and the manufacturer’s protocol is followed for LC3 staining. Invitrogen™ Goat anti-Rabbit IgG (H+L) Cross-Adsorbed Secondary, Antibody, Alexa Fluor™ 488 has been used for secondary staining. Cells were washed and counter stained with DAPI using VECTASHIELD® Antifade Mounting Medium has been done. Microscopy was performed on OLYMPUS IX83 and analyzed as follows: nucleus: DAPI, LC3b: FITC, conidia: Cy5 channel at 20X.

#### siRNA knockdown of *Adipor1*

TriFECTa RNAi Kit was used with Lipofectamine 2000 reagent for transfection. Adherent 10 days old alveolar macrophages were treated with diluted AdipoR1 SiRNA in Lipofectamine (1:1 ratio) at 1nM per 5 x 10^5^ cells concentration for 72 hours. The cells were then quantified for AdipoR1 receptor expression using AdipoR1 RT-PCR and compared with provided positive and negative controls.

### STATISTICAL ANALYSIS

Prism software was used for generation of figures and for statistical analyses (GraphPad). Unpaired *t* tests or ANOVA tests were used to measure statistical significance. Differences between experimental groups that resulted in a *p* value < 0.05 were considered significant.

## Supplemental Information

### Supplemental Methods

### Mice

An additional strain of Adipoq-/- mice was obtained from Dr. Philipp Scherer (University of Texas-Southwestern) and used for survival, fungal burden, histology, flow cytometric analysis of BALF cells, qRT-PCR of selected inflammatory and *Adipor* genes, and ELISA for TNF as described in the Materials and Methods section.

### Quantification of AM AdipoR1 expression using flow cytometry

After ex-vivo AM extraction and culture as mentioned in the methods, AMs were fixed using IC fixation buffer. Fc Block was used to eliminate non-specific Fc-mediated interactions. The cells were then stained at a 1:100 dilution with primary antibody with Adiponectin Receptor 1 Recombinant Rabbit Monoclonal Antibody (SC69-04) followed by staining with Alexa Fluor 488-conjugated goat anti rabbit IgG as the secondary antibody. All the unstained and only primary antibody staining controls were included.

### Quantification of AM *Adipor1* and *Adipor2* receptor gene expression

RNA extraction from AMs was done using Qiagen RNeasy Mini kit, following the manufacturer’s protocol. Quantitative RT-PCR was performed with 20ng of cDNA using AdipoR1 and AdipoR2 forward and reverse primers ordered from IDT Integrated Technologies. Gene Expression Master Mix (ThermoFisher Scientific) was used, with β-actin used for signal normalization.

### RNA sequencing and analysis Sample Preparation

AMs were isolated from APN-/- mice, infected/uninfected for 10 hours (1:9 cells/conidia) and AdipoRon/vehicle treated 24 hours before infection, with cells lysed and RNA isolated as described in Materials and Methods.

### Library preparation and sequencing

Total RNA samples were first evaluated for their quantity and quality using Agilent TapeStation. All the samples were good quality with RIN (RNA Integrity Number) of 9.6-10. One hundred nanograms of total RNA was used for library preparation with the Illumina Stranded mRNA Prep, Ligation kit (lllumina), following the manufacturer’s instruction. Each resulting uniquely dual-indexed library was quantified and quality accessed by Qubit and Agilent TapeStation, and multiple libraries were pooled in equal molarity. The pooled libraries were sequenced with 2×150bp paired-end configuration on an Illumina NovaSeq X PLUS sequencer.

### RNA-seq data analysis

The sequencing reads were first quality checked using FastQC (v.0.11.5, Babraham Bioinformatics, Cambridge, UK) for quality control. The sequence data were then mapped to the mouse reference genome mm10 using the RNA-seq aligner STAR (v.2.7.10a)[52] with the following parameter: “--outSAMmapqUnique 60”. To evaluate quality of the RNA-seq data, the number of reads that fell into different annotated regions (exonic, intronic, splicing junction, intergenic, promoter, UTR, etc.) of the reference genome was assessed using bamutils (from ngsutils v.0.4.17).[53] Uniquely mapped reads were used to quantify the gene level expression employing featureCounts (subread v.2.0.3)[54] with the following parameters: “-s 2 -Q 10”. The data was normalized using TMM (trimmed mean of M values) method. Differential expression analysis was performed using edgeR (v.4.0.1).[54,55] False discovery rate (FDR) was computed from p-values using the Benjamini-Hochberg procedure. Gene Ontology (GO) pathway enrichment analyses were performed with the R package <u>clusterProfiler</u>.[56,57]

---

**Supplemental Video SV1. Time course of uptake and killing in WT and APN-/- AMs.**

Alveolar macrophages were plated and infected as mentioned in the methods. Cells were live imaged by confocal microscopy for T= 8min under 40X water immersion lens. The black arrows indicate the live conidia at T=0 min and the white arrows indicate the conidia at T=8min.

**Fig. S1.**
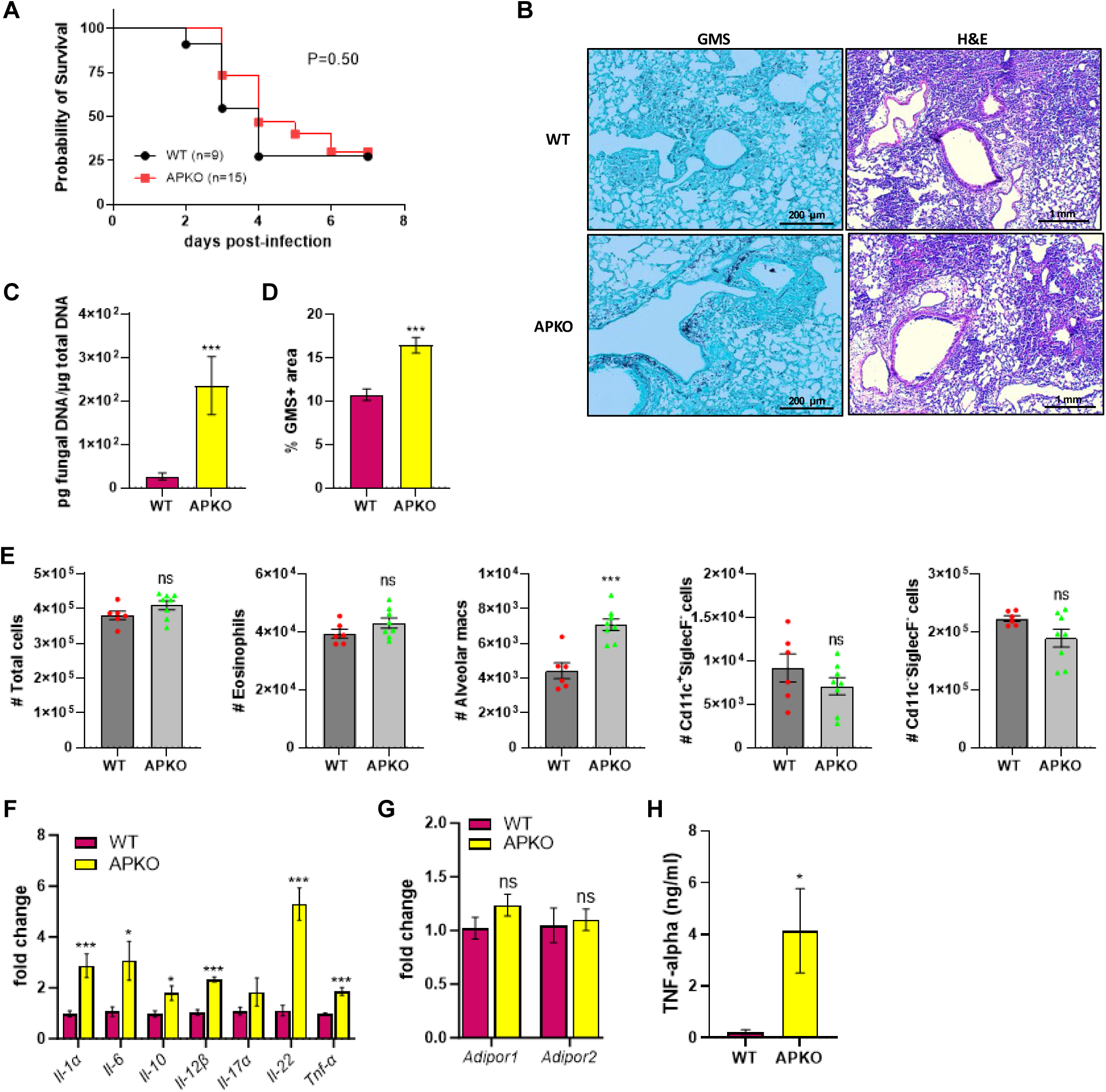
Inflammatory phenotype of APN-/- mice from a second strain. Wild-type (C57BL/6) and APKO (Second *Adipoq*^−/−^ mice from Scherer’s laboratory) were neutrophil depleted and involuntarily aspirated 1 – 1.5 × 10^7^ of conidia as described in Materials and Methods. A. Survival rate. *N* = 9 and 15 mice per group, respectively. B. Representative GMS and H&E lung sections. C. Fungal burden determined by quantitative PCR of fungal DNA from lung homogenates. D. Fungal burden determined by quantification of GMS staining. E. Total number of CD45^+^ cells, eosinophils (CD45^+^Ly6G^−^CD11c^−^SiglecF^+^), AMs (CD45^+^Ly6G^−^CD11c^+^SiglecF^+^), CD11c^+^SiglecF^−^ (CD45^+^Ly6G^−^ CD11c^+^SiglecF^−^), and CD11c^−^SiglecF^−^ (CD45^+^Ly6G^−^CD11c^−^SiglecF^−^) cells isolated from the mice with IA as determined by flow cytometry. *N* = 4-6 mice per group. F. qRT-PCR analysis for mRNA expression of the indicated cytokines. G. qRT-PCR analysis for mRNA expression of *Adipor1* and *Adipor2*. H. TNFα secretion in BALF quantified at the protein level by ELISA. Data are a summary of two to three independently performed experiments. \**p* < 0.05, \*\**p* < 0.01, \*\*\**p* < 0.001.

**Fig. S2.**
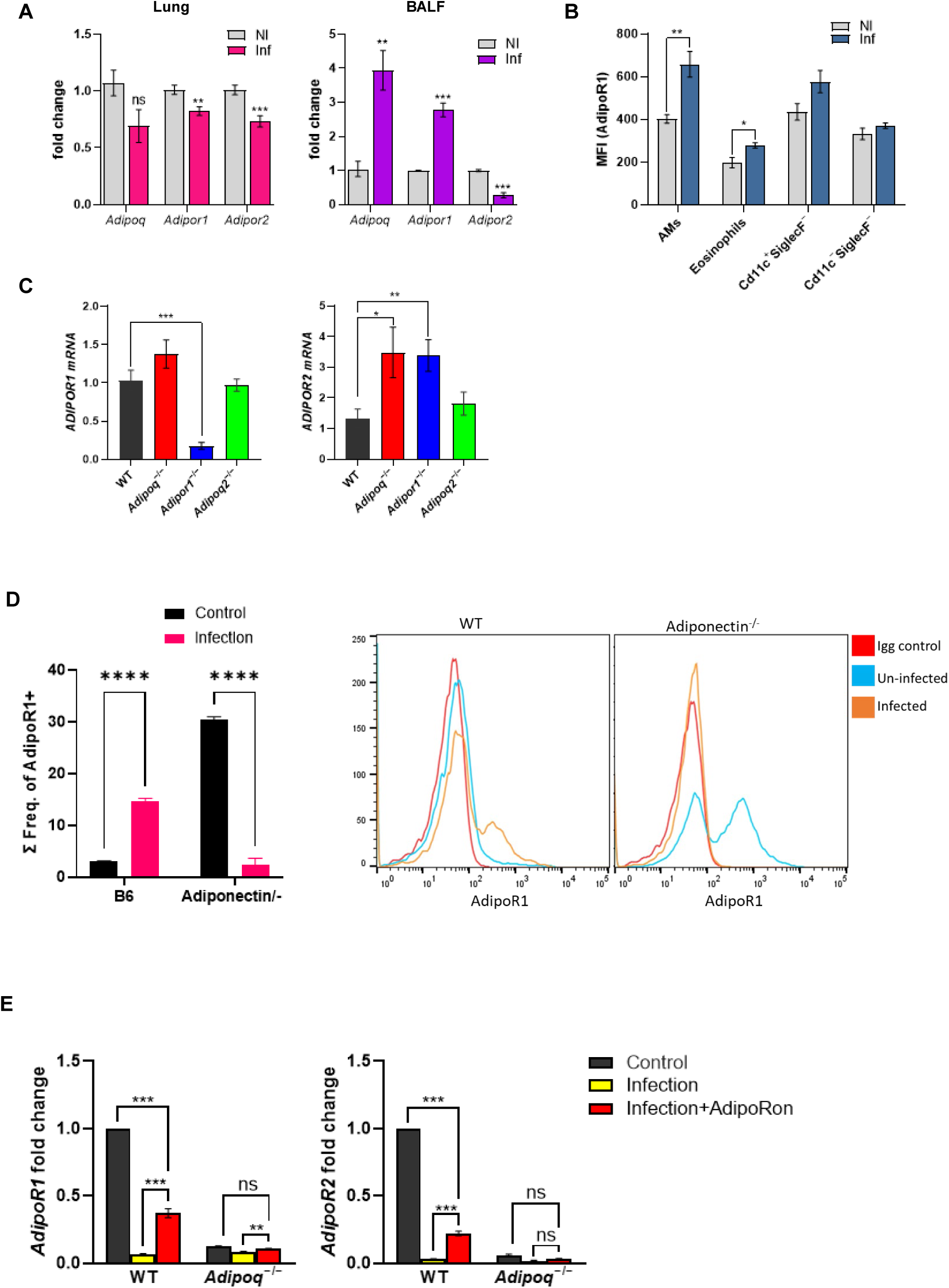
AdipoRs expression in the Lung and BALF in APN pathway-deficient mice with invasive aspergillosis. A. Expression of *Adipoq* and *Adipor* genes in lung homogenates and BALF cells from non-infected (NI) and conidia-infected WT mice. Lung and BALF were collected from the mice at 3 dpi. B. Summary of median fluorescence intensities of AdipoR1 staining on AMs (CD45^+^Ly6G^−^CD11c^+^SiglecF^+^), eosinophils (CD45^+^Ly6G^−^CD11c^−^SiglecF^+^), CD11c^+^SiglecF^−^ (CD45^+^Ly6G^−^ CD11c^+^SiglecF^−^), and CD11c^−^SiglecF^−^ (CD45^+^Ly6G^−^CD11c^−^SiglecF^−^) cells from non-infected (NI) and conidia-infected WT mice. C. qRT-PCR analysis for mRNA expression of *Adipor1* and *Adipor2* in lung homogenates. Wild-type (C57BL/6), *Adipoq*^−/−^, *AdipoR1*^−/−^, and *AdipoR2*^−/−^mice were neutrophil depleted and involuntarily aspirated *A. fumigatus* conidia. D. Flow cytometry staining of AdipoR1. Frequency of AdipoR1+ in ex-vivo cultured AMs in WT and *Adipoq*^−/−^ mice. The histogram represents the AdipoR1 peak relative to IgG control in infected vs uninfected. E. qRT-PCR analysis for mRNA expression of *Adipor1* and *Adipor2* from the ex-vivo cultured AMs. The AMs were challenged with AF-293 conidia with or without AdipoRon treatment. Data are a summary of two to three independently performed experiments. \**p* < 0.05, \*\**p* < 0.01, \*\*\**p* < 0.001.

**Fig. S3.**
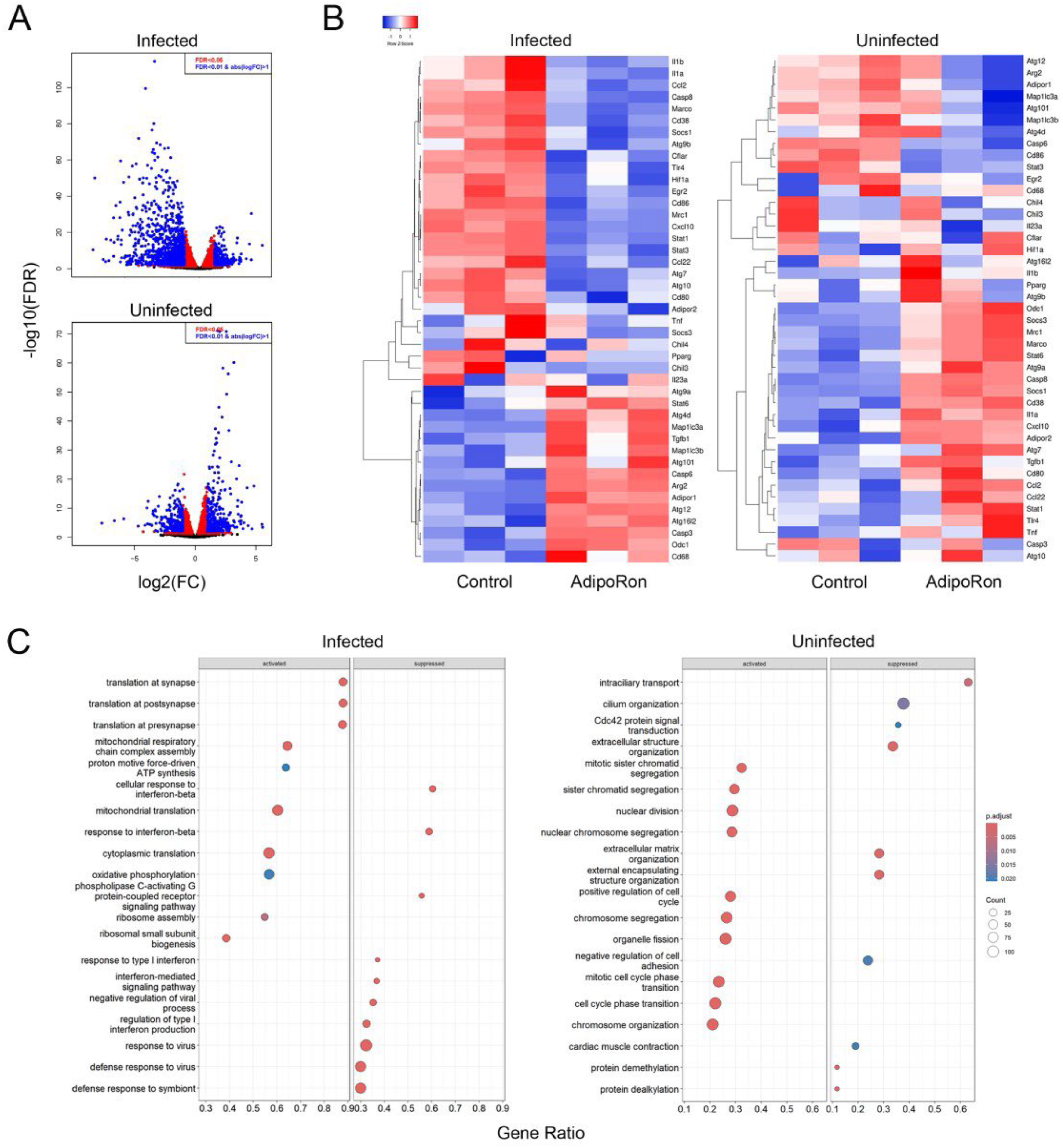
Gene expression in AdipoRon treated/untreated and infected/ uninfected *Adipoq^−/−^* AMs by RNAseq analysis. AMs were infected with swollen AF293 conidia with 1:9 cells/conidia for 10 hours, or left uninfected, with or without AdipoRon treatment, followed by RNA extraction for RNAseq analysis (N=3/group). A. Volcano plot depicting relative changes in gene expression of AdipoRon-treatment in APN-/- AMS, infected (top) or uninfected (bottom). B. Heat map representation of the genes with highest differential expression in infected (left) and uninfected (right) APN-/- AMs. C. GSEA-GO analysis of gene pathways that are both differentially expressed by AdipoRon treatment in infected (left) and uninfected (right) APN-/- AMs.

